# Biology and rearing of the cogongrass gall midge, *Orseolia javanica* Kieffer & Docters van Leeuwen-Reijnvaan (Diptera: Cecidomyiidae)

**DOI:** 10.1101/2020.02.27.966499

**Authors:** Purnama Hidayat, Arini, Dwi Guntoro, Keiji Takasu, William A. Overholt

## Abstract

*Imperata cylindrica* (L.) Beauv. (Poaceae) is one of the most harmful weeds in the world because of its ability to spread and form high density, monospecific stands that exclude other vegetation. The cogongrass gall midge, *Orseolia javanica* Kieffer & Docters van Leeuwen-Reijnvaan (Diptera: Cecidomyiidae), is a stem galling insect that is only known to develop in cogongrass and has only been found on the island of Java in Indonesia. The midge attacks very young shoots, which stimulates abnormal growth, resulting in the formation of a purplish, elongate stem gall tappered to a point at the apical end. The aim of the current research was to describe the biology of the midge and develop a rearing method. *Orseolia javanica* completed its life cycle in 12-38 days with average egg, larval, and pupal periodes of 4.0 ± 0.0, 13.5 ± 3.8, and 8.6 ± 6.6 days (mean ± SD), respectively. Mated female, unmated female, and male longevities were 1.7 ± 0.47, 1.2 ± 0.41, and 1.0 ± 0.00 days (mean ± SD). Galls began to appear 29 days after larval infestation, and stem death coincided with emergence of the adult midge. The midge may have potential for biological control of cogongrass if future studies confirm a restricted host range.

---

Cogongrass, *Imperata cylindrica* (L.) Beauv. (Poaceae), is a serious weed in many areas of the world including Asia (Brook 1989; Garrity *et al.* 1997), West Africa (Chikoye & Ekeleme 2001) and the southeastern United States (McDonald 1996). In Asia, the largest infested area occurs in Indonesia (Garrity *et al.* 1997). The native range of cogongrass is vast and thought to include Africa, southern Europe, Asia and northern Australia (Hubbard *et al.* 1944), while populations in the southeastern USA are exotic and highly invasive (Burrell *et al.* 2015). There has been some speculation that the center of origin of cogongrass is East Africa because of the high diversity of plant pathogens (Ivens 1983), lack of weediness and high genetic diversity (Overholt *et al.* 2016). Cogongrass spreads rapidly by seeds and rhizomes, often forming monospecific stands that exclude other vegetation, resulting in both ecological and economic damage (Brook 1989; McDonald 1996). Moreover, cogongrass increases the frequency and intensity of wildfires (Jose *et al.* 2002).

In 2013, the University of Florida initiated exploration for natural enemies of cogongrass in several countries in Africa and Asia with the objective of identifying insects that may have value for introduction into the USA as biological control agents (Overholt *et al.* 2016). One of the insects encountered during surveys in Indonesia was *Orseolia javanica* Kieffer & Docters van Leeuwen-Reijnvaan (Diptera: Cecidomyiidae). The midge is a gall-forming insect that is only known to develop in cogongrass (Mangoendihardjo 1980; Soenarjo 1986). Early instar larvae of *O. javanica* colonize young cogongrass shoots which stimulates the formation of elongate, purplish stem galls, tapered to a point at the apical end (Mangoendihardjo 1980). Based on field observations, the gall midge can be highly damaging and may have potential for biological control for cogongrass in the southeastern USA and elsewhere (Overholt *et al.* 2016).

The cogongrass gall midge has only been found in West and Central Java in Indonesia (Mangoendihardjo 1980). The population dynamics of the gall midge and its parasitoids in Cianjur, West Java were recently reported by Aviansyah (2016), who found that parasitoids emerged from 40-60% of galled stems. Similarly, Buhl *et al.* (2016) reported that three species of parasitoids emerged from 64% of galled stems collected in June and July 2015. In addition, Gumilang (2016) provided information about the distribution of the gall midge in Bogor and Cianjur Districts in West Java and Magelang and Salatiga Districts in Central Java.

The objective of the current study was to provide basic biological information about the midge, which is essential for developing a laboratory rearing method. The ability to rear the midge is a critical first step towards the initiation of host ranges studies required to evaluate the potential safety of the midge as a classical biological control agent.

## MATERIALS AND METHODS

### Plants

Cogongrass used in the laboratory studies was collected at Leuwikopo, Bogor Agricultural University. Plants were excised from the soil and planted in 25 plastic trays (40 cm × 27 cm × 15 cm, L, W, H) with each pot receiving 20 stems. Trays were held outdoors inside screen cages (240 cm × 120 cm × 120 cm, L, W, H) for one month prior to initiation of studies. The one month period insured that stems were not naturally infested with gall midges. Studies were conducted at the Laboratory of Insect and Biosystematics, Department of Plant Protection, Faculty of Agriculture, Bogor Agricultural University.

### Insects

Galled cogongrass stems (45-130) were collected on eight sampling occasions between August and November 2016 from bunds bordering rice fields near the village of Desa Cibereum in Cugenang District, Cianjur Regency, West Java. Desa Cibereum is located at - 6.7914 latitude, 107.0687 longitude, at 852 m above sea level. In total, 832 galled stems were collected on the eight sampling occasions. Galled stems, along with attached rhizome, were removed from the soil and transported to the laboratory in coolers. Galls were then placed individually in plastic tubes and held at ambient laboratory conditions (27.3 ± 0.21 °C, 68 ± 0.81 RH) for emergence of adult midges. The size of the tubes varied from 5 cm × 8 cm to 5.5 cm × 30 cm (D, H) depending on the length of the gall. Images of the midge’s life stages and measurements were made using a Leica M205C stereo microscope attached to a Leica DFC 450 camera and processed using Leica Application Suite Version 4.4.0 software. All estimates of central tendencies are presented as means ± standard deviation. Voucher specimens of adults and immature stages are maintained in the insect collection at the Laboratory of Insect Biosystematics, Department of Plant Protection, Faculty of Agriculture, Bogor Agricultural University.

### Oviposition, Egg Morphology and Development

Adult males and females which emerged in tubes were placed together in the laboratory at ambient conditions in varying numbers in a plastic box (17 cm × 12 cm × 12 cm, L, W, H) that had a moist tissue paper on the bottom. A window (13 cm × 7 cm, L, W) was cut in the top of the box and covered with fine mesh cloth to allow ventilation. Eggs were laid by females on the bottom and side of the plastic box and collected after adults died. Eggs (20 to 50) were transferred to moist tissue paper in petri dishes (2 cm × 9 cm, D, H) using a fine brush. The length and width of eggs were measured and then observed daily until eclosion. Developmental time and the percentage of eggs that hatched were determined from a sample of 150 eggs.

### Larval and Pupal Morphology and Development

Newly hatched larvae were transferred using a fine brush to the cogongrass stems which had been planted in trays, with each stem receiving one larva. Stems were trimmed to 2 cm above the soil surface prior to infestation. A total of 400 larvae were placed on cogongrass stems. Stems were held outdoors in large screen cages. Starting five days after infestation, 5-10 cogongrass stems were randomly selected and dissected every day to access larval colonization and growth of the larvae. In total, 350 stems were observed over a period of 57 days. Dissections were performed under a stereo microscope using a scalpel and a micro needle. Larvae and pupae were preserved in 70% ethanol in Eppendorf tubes (1.5 ml). A subset of 20 larvae was slide mounted to measure their length as the distance from the sternal spatula to the front of the head, and width at the widest point. Slides were prepared using a method described in Watson (2007), modified as follows: the posterior part of the larvae was pierced with a micro needle and then boiled in 95% ethanol for 3 minutes. After 3 minutes, 10% KOH was added until the larvae appeared clear. The body contents were then removed under a stereo microscope using a micro needle. The larval skins were washed twice in distilled water and then dehydrated for five minutes each in a series of ethanol concentrations along with 1-2 drops of acid fuchsin, starting with 50% and proceeding to 80%, 95%, and finally to 100%. After that, the larvae were immersed in clove oil and slide mounted in Canada Balsam.

### Adult Behavior, Morphology, Fecundity and Longevity

The behavior of adults was observed from the time that they emerged until mating. A subset of 38 field collected adults (19 males, 19 females) were killed in 70% ethanol and measured to determine body length, wingspan (distal end of left wing to distal end of right wing), and width at the widest point of the thorax. Sizes of males and females were compared using a two-sample *t*-test. Adult longevity was determined from a sample of 13 males, 11 mated females, and 30 unmated females. Fecundity was determined by counting the number of eggs laid by 10 mated and 10 unmated females from emergence until death.

### Gall Size

The length and diameter of galls from which males and females (7 each) had emerged were measured and compared using a two-sample *t*-test. The galls were collected from the field in Cianjur, West Java.

## RESULTS

### Oviposition, Egg Morphology and Development

From the 832 galls collected, 55 females and 44 males emerged. Eggs were laid singly or in groups of 2-18, side by side and longitudinally parallel on the wall of the plastic box or on the moist tissue paper on the bottom of the box. Eggs eclosed on the fourth days after oviposition. Eggs that failed to hatch became transparent or moldy.

Eggs were oval shaped with a length of 0.53 ± 0.004 mm and a width of 0.14 ± 0.001 mm (Table 1). All freshly laid eggs were yellowish white. On the second day after oviposition, eggs from mated females became red at both the apical and distal ends. On the third day, the red color at both ends of the eggs faded while the lateral edges reddened, with the color becoming increasingly apparent as the eggs matured on the fourth day (Figure 1a-d). Eggs from unmated females remained transparent and did not undergo any color change.

**Table 1.** Sizes and developmental durations (means ± SD) of immature stages of cogongrass gall midge.

| Stages | Length<br>(mm) | Width<br>(mm) | Length of<br>sternal<br>spatula<br>(mm) | Width of<br>head<br>(mm) | Duration<br>(days) | N |
| --- | --- | --- | --- | --- | --- | --- |
| Egg | 0.53±0.004 | 0.14± 0.001 | - | - | 4.0±0.0 | 133 |
| Larva |  |  |  |  |  |  |
| First instar | 0.59±0.100 | 0.19±0.010 | - | 0.05±0.01 | 7.1±1.4 | 10 |
| Second instar | 1.18±0.400 | 0.43±0.150 | - | 0.09±0.02 | 2.7±1.5 | 12 |
| Third instar | 3.30±1.500 | 1.00±0.350 | 0.09±0.02 | 0.08±0.01 | 9.8±4.7 | 32 |
| Pupa | 5.89±0.700 | 1.61±0.250 | - | - | 8.5±6.6 | 30 |

**Figure 1.**
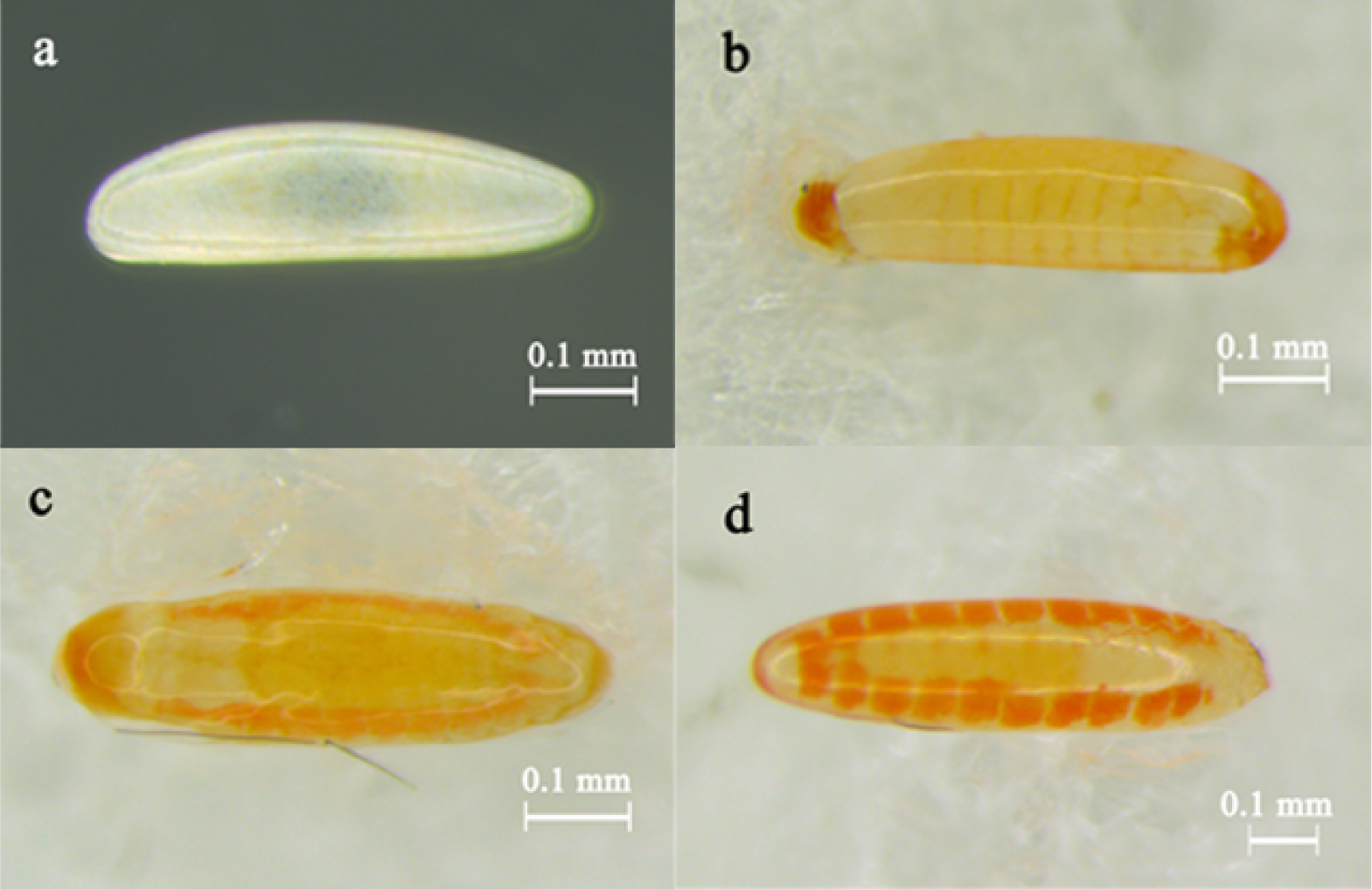
One (a), two (b), three (c) and 4 days (d) old eggs of *O. javanica*.

### Larval and Pupal Morphology and Development

Of the 350 stems that were inoculated with neonate larvae and dissected, 27% were successfully infested by gall midges and exhibited signs of deformation, 21% of stems died due to decay at the roots and the remaining 52% of stems were not colonized. Abnormal stem growth became evident 10 days after inoculation as an enlargening of the stem at the growing point. During early dissections (< 10 days after inoculation), some stems were found to be infested with two larvae, but later dissections determined that only one larva survived to pupation. Developmental time and the average length and width of the body, length of the sternal spatula and width of the head are shown in Table 1. Newly hatched larvae were 0.49 ± 0.03 mm long and 0.17 ± 0.006 mm wide, reddish to orange, transparent, and had a black eye spot (Figure 2a). The second instar larva was orange to white with a black eye spot (Figure 2b). The dorsal side of second instar larvae was convex whereas the ventral side tended to be flat. Third instar larvae were white to yellowish and their bodies were firmer than those of second instars. The segments of third instars were more apparent than earlier instars and the sternal spatula became visible (Figure 2c).

**Figure 2.**
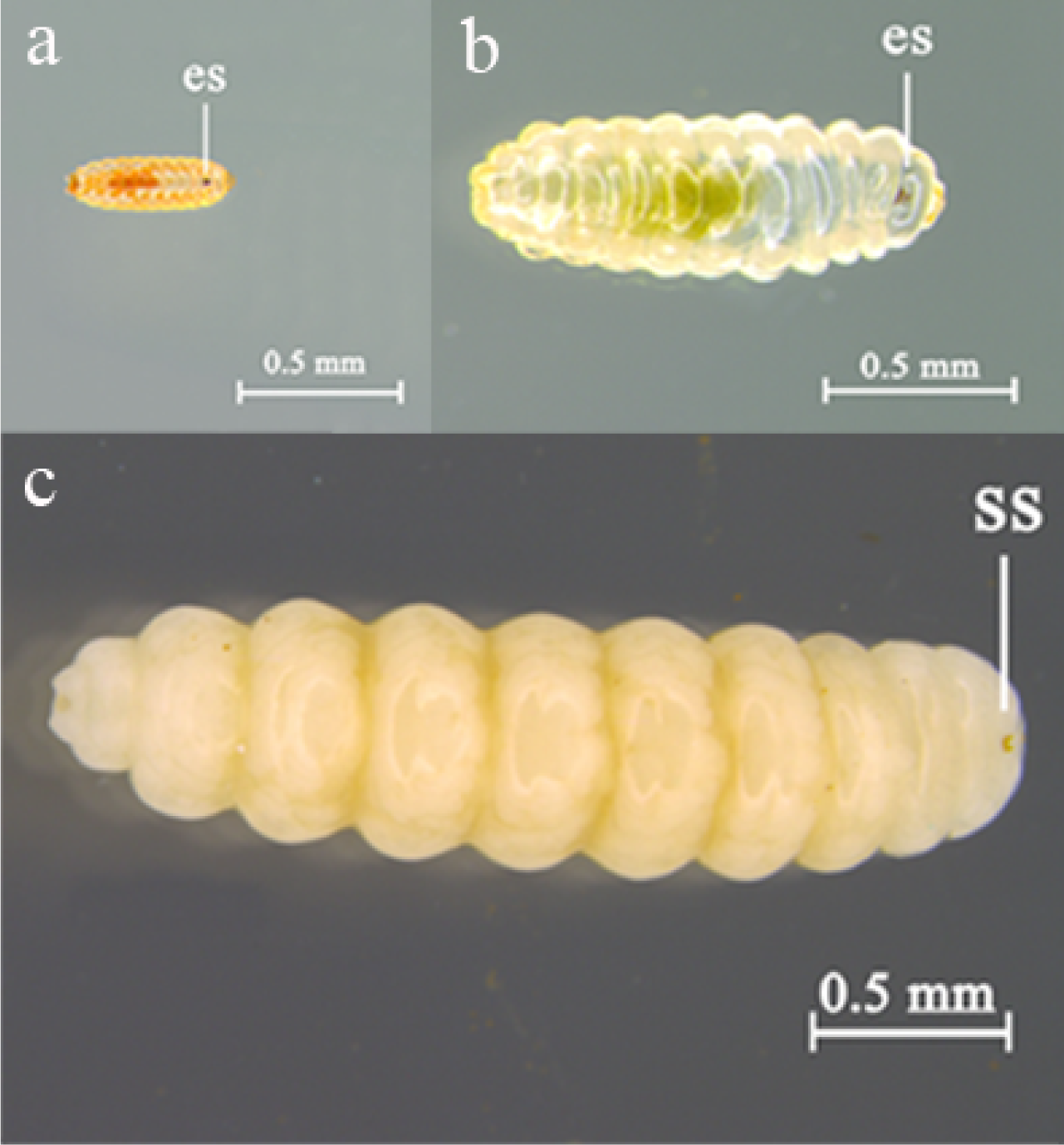
Neonate (a), 11 days old (b), and 21 days old larvae of *O. javanica*, es, eye spot; ss, sternal spatula.

Pupae of cogongrass gall midge were first white, and then darkened as they aged, becoming orange, light brown and dark brown with the eyes, legs and antennae black (Figure 3a-c). The length and width were 5.89 ± 0.72 mm and 1.61 ± 0.25 mm, respectively. The mean developmental time of pupae was 8.5 ± 6.6 days (Table 1).

**Figure 3.**
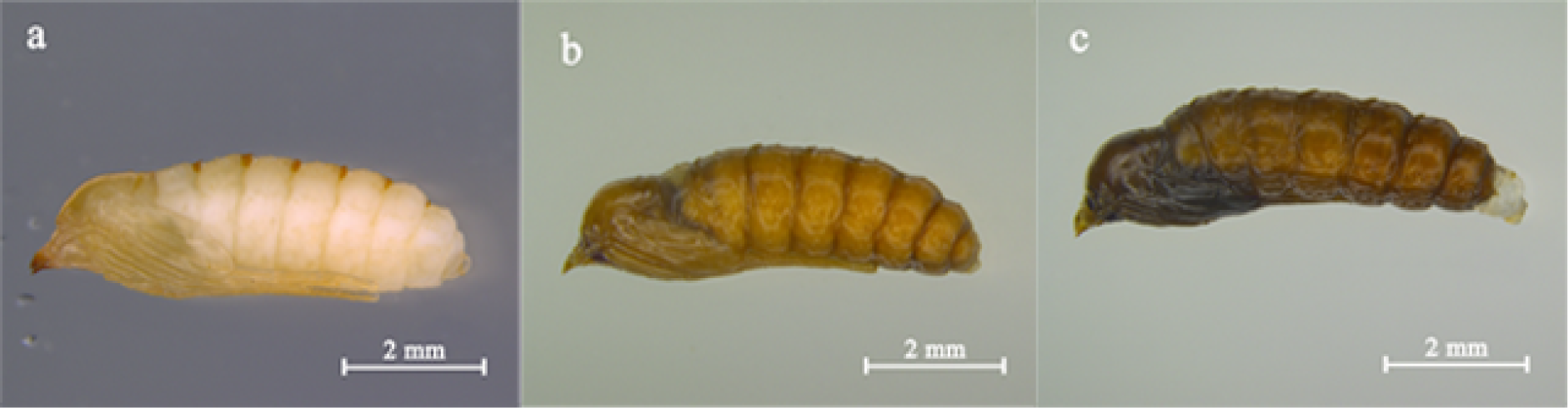
Pupae of *O. javanica* at 7 (a), 15 (b), and 25 days (c) after pupation.

### Adult Behavior, Morphology, Fecundity and Longevity

Adult emergence occurred predominantly in the morning, but some individuals emerged in the afternoon and evening. After emerging from galls, adults remained on the outside of the gall for several minutes and then males flew off, while females tended to remain on the gall from which they emerged or on the soil nearby. Adults were covered with fine hairs on all body surfaces. Males were slender with black bodies and much smaller than females. Females were much broader than males and had a brown abdomen (Table 2, Figure 4a-b). Lifetime fecundity of mated females was 512.1 ± 194.4 eggs, while unmated female laid 153.3 ± 86.3 eggs (Figure 5a-b). Mated female, unmated female, and male longevities were 1.7 ± 0.47, 1.2 ± 0.41, and 1.0 ± 0.00 days (mean ± SD).

**Table 2.** Length, width and wingspan (mean ± SD) of adult male and female cogongrass gall midge.

| Parameter | Male (mm)<sup>a</sup> | Female (mm)<sup>a</sup> | *t* | df | P |
| --- | --- | --- | --- | --- | --- |
| Body length (mm) | 2.95±0.27a | 5.65±1.12b | 10.3 | 36 | < 0.00001 |
| Body width (mm) | 0.58±0.06a | 1.16±0.15b | 15.8 | 36 | < 0.00001 |
| Wingspan (mm) | 6.63±1.19a | 9.76±0.49b | 10.5 | 36 | < 0.00001 |

<sup>a</sup> Means in a column followed by different lowercase letters are significantly different ( $P \leq$ 0.05; ANOVA and LSD test).

**Figure 4.**
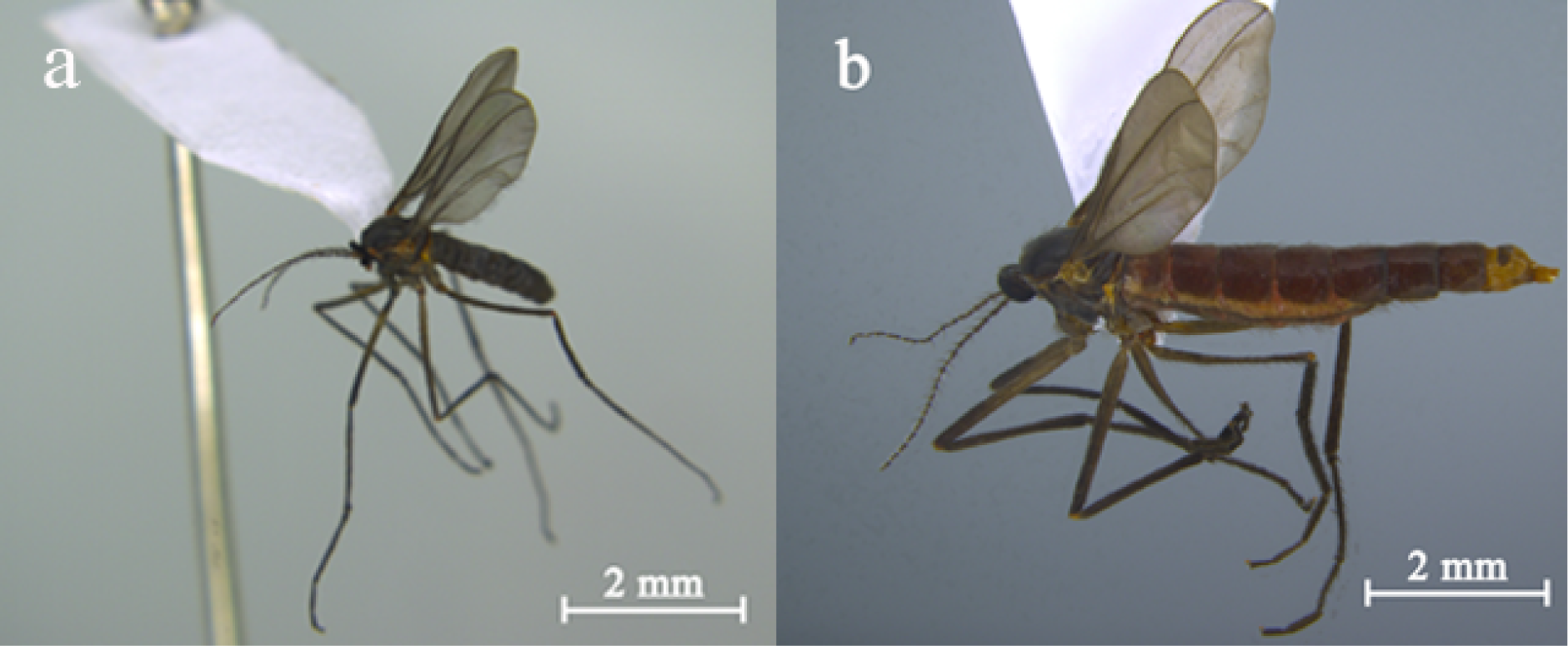
Adult male (a) and female (b) cogongrass gall midges.

**Figure 5.**
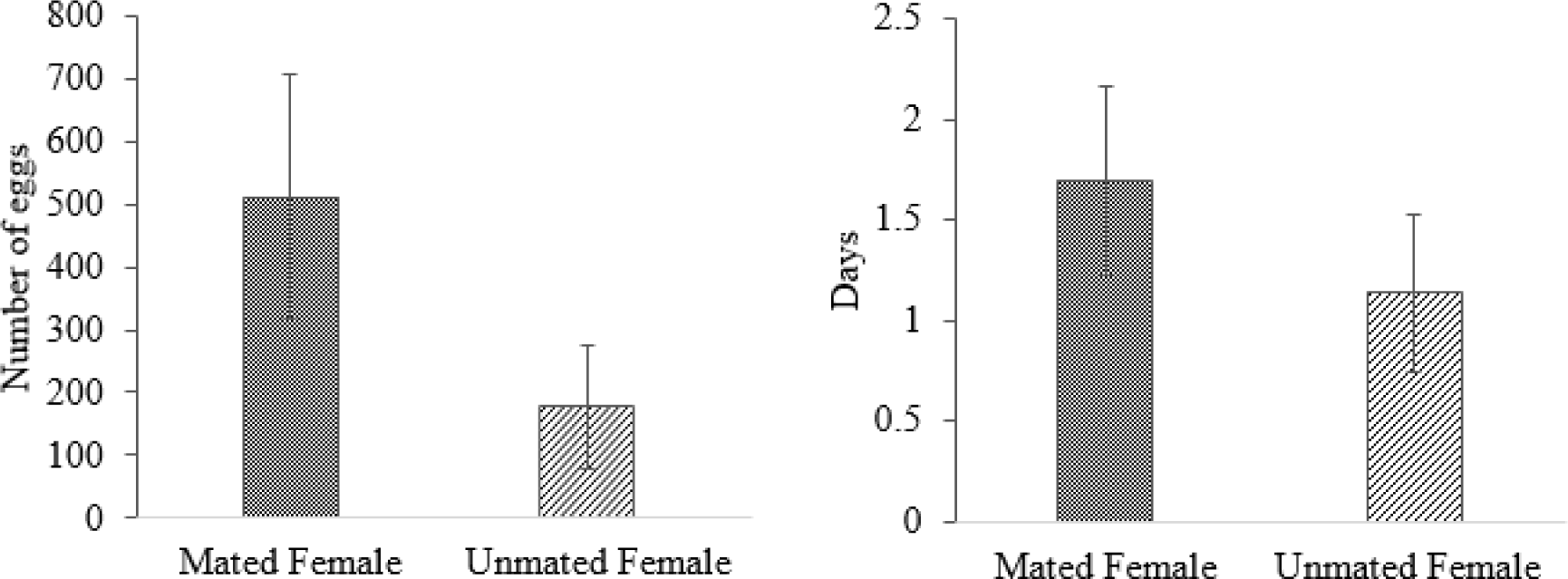
Fecundity (a) and longevity (b) (mean ± SD) of adult female cogongrass gall midges.

### Gall size

Gall morphology differed depending on whether a male or female emerged. Galls from which males emerged were longer (male: 125 ± 45 mm, female: 80 ± 23 mm; *t* = 2.4, df = 12, P = 0.02)) and thinner (male: 2.5 ± 2 mm, female: 3.2 ± 2 mm; *t* = 8.9, df = 12, P < 0.00001) than galls from which females emerged.

## DISCUSSION

Developmental time of the midge (range = 12-38 days, mean = 25.4 ± 7.6 days) was surprisingly variable, and much less than that found by Soenarjo (1986) of 35 to 49 days. The variability in developmental times of larvae and pupae in our study was likely due to fluctuations in ambient outdoor temperatures where the infested stems were maintained. Mangoendihardjo (1980) also reported much longer developmental times than those in our study (33 days for females and 35 days for males) but provided no measure of variability. The developmental times we found are similar to those reported for other *Orseolia* spp. *Orseolia oryzae* developed in 15 days (Jagadeesha Kumar 2009) or 19-23 days (Rajamani et al. 2004). *Orseolia oryzivora* was found to complete development in 26 days (Umeh & Joshi 1993).

Unfortunately, due to the lack of detail provided in Mangoenhihardio (1980), it is difficult to compare the success of the two rearing methods. The proportion of stems that were successfully colonized by the midge larvae in our study (27%) was low. Reasons for the relatively low success in colonization are unknown but could be due to injury to the delicate neonate larvae during transfer to stems, or perhaps differences in the susceptibility of stems due to their age, size or physiological state. Mangoendihardjo (1980) also reported a low success rate of stem colonization with only 25% of stems exposed to ovipositing females successfully colonized. Surprisingly, he also found that the majority (98%) of *O. javanica* eggs were laid on the soil surface, with only 2% laid on the plant. If this is representative of what occurs in nature, we suspect that there is very high mortality of larvae during their search for suitable stems due to predation and desiccation.

*Orseolia javanica* has three larval instars, as had been found for other *Orseolia* spp., including *O. oryzae* (Perera & Fernando 1970) and *O. oryzivora* (Ogah et al. 2010). Similar to our findings, first and second instars of both *O. oryzae* and *O. oryzivora* had visible eyespots, and the third instar was characterized by the presence of a sternal spatula (Perera & Fernando, 1970; Ogah et al. 2010).

Biological control of weeds utilizes highly host specific insect herbivores to regulate exotic weeds in their areas of invasion. A narrow host range is required to insure that introduced biological control agents will have little or no negative impacts to native flora or cultivated crops after release. Additionally, biological control scientists are increasingly encouraged to conduct studies on the potential impact of candidate biological control agents to the target plant in order to avoid the introduction ineffective agents (McClay & Balciunas 2005). In order to delineate the physiological host range of candidate biological control agents and conduct impact studies, effective rearing methods are required. Our study provides basic biological information and describes a rearing method for *O. javanica*, a potential biological control agent of cogongrass in the southeastern USA (Overholt *et al.* 2016).

Limited host range testing of *O. javanica* suggests that it may have the requisite host specificity for release as a biological control agent in USA. Mangoendihardjo (1980) exposed cogongrass, three varieties of cultivated rice, two wild rice species of questionable identification (published as *Oryza fatua*, which is considered a synonym of *O. rufipogon* Griff. and *O. perennis* which is of doubtful taxonomic status and may also be a synonym of *O. rufipogon* (Terrell *et al.* 2000), sorghum (*Sorghum bicolor* (L.*) Moench), maize (Zea mays* L.) and two wild grasses (*Paspalum conjugatum* Berguis, *Pennisetum polystachyon* (L.) Schult.) to the midge and it only completed development in cogongrass. Narrow host ranges of other *Orseolia* spp. also point towards a possible high specificity of *O. javanica*. The African species, *O. bonzii* Harris is only known to develop in *Paspalum scobiculatum* (L.), and *O. nwanzei* Harris & Nwilene only in *Eragostris atrovirens* (Desf.) Trin. ex. Steud (Nwilene *et al.* 2006), while the Africa rice gall midge, *O. oryzivora*, only completes development on cultivated and wild *Oryza* spp. (Williams *et al.* 1999). The Asian rice gall midge, *O. oryzae* was found only in cultivated rice during a survey of 10 wild grasses and wild *Oryza* spp.in the state of Karnataka, India (Kumar et al. 2009), while another study in India reported the midge from two wild grasses and three *Oryza* spp. (Rajamani *et al.* 2004). Twenty-four species of oriental *Orseolia* are described, and the majority, including *O. javanica*, have only been collected from one grass host (Gagné & Jaschhof 2017).

In summary, our studies provide basic biological information, and describe a rearing method that can be used to initiate and maintain laboratory colonies of *O. javanica* so that host range and impact studies can be conducted in order to determine whether the midge may be considered for release as a biological control agent of cogongrass.

## ACKNOWLEDGMENTS

This research was partly funded by the Thesis and Dissertation Grant from The Indonesian Endowment Fund for Education (LPDP), Ministry of Finance, Republic of Indonesia and the Florida Agricultural Experiment Station, Florida Fish and Wildlife Conservation Commission, the Florida Department of Agriculture and Consumer Services.

## LITERATURE CITED

Aviansyah, I. 2016. Population and parasitoid of the congongrass gall midge *Orseolia javanica* Kieffer & Van Leeuwen-Reijnvaan (Diptera: Cecidomyiidae) in Cianjur Regency] [BSc Thesis]. Institut Pertanian Bogor, Bogor.

Brook, R.M. 1989. Review of literature on *Imperata cylindrica* (L.) Raeuschel with particular reference to South East Asia. Tropical Pest Management 35: 12–25.

Buhl, P.N. and Hidayat, P. 2016. A new species of *Platygaster* (Hymenoptera: Platygastridae) reared from *Orseolia javanica* (Diptera: Cecidomyiidae) on cogongrass, *Imperata cylindrica* (Poaceae). International Journal of Environmental Studies 73:25–31.

Burrell, M., Pepper, A.E., Hodnett, G., Goolsby, J.A., Overholt, W.A., Racelis, A.E., Diaz, R., and Klein, P.E. 2015. Exploring origins, invasion history and genetic diversity of *Imperata cylindrica* (L.) P. Beauv. (Cogongrass) in the United States using genotyping by sequencing. Molecular Ecology 24:2177–2193.

Chikoye, D. and Ekeleme, F. 2001 Weed flora and soil seedbanks in fields dominated by *Imperata cylindrica* in the moist savannah of West Africa. Weed Research 41:475–490.

Gagné, R.J. and Jaschhof, M. (2017) *A Catalog of the Cecidomyiidae (Diptera) of the World*, 4^th^ edn. Systematic Entomology Laboratory, Agricultural Research Service, U.S. Department of Agriculture, Washington.

Garrity, D., Soekardiz, M., Van Noordwijk, M., De La Cruz, R., Pathak, P., Gunasenas, H., and Van So, N. 1997. The *Imperata* grasslands of tropical Asia: area, distribution, and typology. Agroforestry Systems 36:3–29.

Gumilang, B.L. 2016 Distribution and population level of the congongrass gall midge, *Orseolia javanica* Kieffer & van Leeuwen-Reijnvaan (Diptera: Cecidomyiidae) [BSc Thesis]. Institut Pertanian Bogor, Bogor.

Hubbard, C.E., Whyte, R.O., Brown, D., and Gray, A.P. 1944. *Imperata cylindrica*: taxonomy, distribution, economic significance and control. Imperial Agricultural Bureaux Joint Publication 7:1–63.

Ivens, G.W. 1983. The natural control of *Imperata cylindrica*: Nigeria and northern Thailand. Mountain Research and Development 3:372–377.

Jagadeesha Kumar, B.D., Chakravarthy, A.K., Doddabasappa, B., and Basavaraju, B.S. 2009. Biology of the rice gall midge, *Orseolia oryzae* (Wood-Mason) in southern Karnataka. Karnataka Journal of Agricultural Science 22:535–537.

Jose, S., Cox, J., Miller, D.L., and Shilling, D.G. 2002. Alien plant invasions: the story of cogongrass in southeastern forests. Journal of Forestry 100:41–44.

Kumar, B.D.J., Charkravarthy, AK, Doddabasappa, B., and Basavaraju, B.S. 2009. Biology of the rice gall midge, *Orseolia oryzae* (Wood-Mason) in southern Karnataka. Karnataka J. Agricultural Science 22: 535–537.

Mangoendihardjo, S. 1980. Some notes on the natural enemies of alang-alang (*Imperata cylindrica* (L.) Beauv.) in Java, pp. 47–55. In Proceedings of BIOTROP Workshop on Alang-alang. Jul 27-29 1976, Bogor, Indonesia (Edited by Soewardi). Seameo Biotrop, Bogor.

McClay, A.S. and Balciunas, J.K. 2005. The role of pre-release efficacy assessment in selecting classical biological control agents for weeds—applying the Anna Karenina principle. Biological Control 35:197–207.

McDonald, S.K., Shilling, D.G., Okoli, C.A.N., Bewick, T.A., Gordon, D., Hall, D., and Smith, R. 1996. Population dynamics of cogongrass, Imperata cylindrica, pp. 156. In Proceedings of the Southern Weed Science Society.

Nwilene, F.E., Nwanze, K.F., and Olahidievlbie, O. 2006. *African Rice Gall Midge, Biology, Ecology and Control*. Field Guide and Technical Manual. African Rice Centre (WARDA) Cotonou, Benin. 24 pp.

Ogah, E., Odebiyi, J., Ewete, F., Omoloye, A., and Nwilene, F. 2010. Biology of the African rice gall midge *Orseolia oryzivora* (Diptera: Cecidomyiidae) and its incidence on wet-season rice in Nigeria. International Journal of Tropical Insect Science 30:32–39.

Overholt, W.A., Hidayat, P., Le Ru, B., Takasu, K., Goolsby, J.A., Racelis, A., Burrell, A.M., Amalin, D., Agum, W., Njaku, M., Pallangyo, B., Klein, P.E., and Cuda, J.P. 2016. Potential biological control agents for management of cogongrass (Cyperales: Poaceae) in the southeastern USA. Florida Entomologist 99:734–739.

Perera, N. and Fernando, H.E. 1970. Infestation of young rice plants by the rice gall midge, *Pachydiplosis oryzae* (Wood-Mason) (Diptera: Cecidomyiidae), with special reference to shoot morphogenesis. Bulletin of Entomological Research 59:605–613.

Rajamani, S., Pasalu, I.C., Mathur, K.C., and Sain, M. 2004. Biology and ecology of rice gall midge, *Orseolia oryzae* (Wood-Mason), pp. 7-16. In *New Approaches to Gall Midge Resistance in Rice*. Proceedings of the International Workshop. 22-24 November 1998, Hyderabad, India (Edited by J Bennett, JS Bentur, IC Pasalu, K Krishnaiah). International Rice Research Institute, Hyderabad, India.

Soenarjo, E. 1986. Alang-alang gall midge potential as an alternate host for parasitoids. International Rice Research Newsletter 11:22–23.

Terrell, E.E. and Duvall, M.R. 2000. Oryza. Catalogue of New World Grasses (Poaceae): I. Subfamilies Anomochlooideae, Bambusoideae, Ehrhartoideae, and Pharoideae. Contributions from the United States National Herbarium 39:89–92.

Umeh, E.D.N. and Joshi, R.C. 1993. Aspects of the biology, ecology and natural biological control of the African rice gall midge, *Orseolia oryzivora* Harris and Gagné (Diptera: Cecidomyiidae) in central and southeast Nigeria. Journal of Applied Entomology 116:391–398.

Watson, G.W. 2007. Identification of whiteflies (Hemiptera: Aleyrodidae), pp. 16-26. In APEC Re-entry Workshop on Whiteflies and Mealybugs. 16–26 April 2007, Kuala Lumpur, Malaysia.

Williams, C.T., Okhidievbie, O., Harris, K.M., and Ukwungwu, M.N. 1999. The host range, annual cycle and parasitoids of the African rice gall midge *Orseolia oryzivora* (Diptera: Cecidomyiidae). Bulletin of Entomological Research 89:589–597.

